# Regional analysis of spatiotemporal trends in Colorado potato beetle abundance

**DOI:** 10.64898/2025.12.31.697201

**Authors:** Nursultan Azhimuratov, Russell L. Groves, Juan Francisco Mandujano Reyes, Jun Zhu, Sean D. Schoville

## Abstract

Crop pests can significantly damage crops, cause economic loss, and reduce the sustainability of agroecosystems. New ecoinformatic approaches are needed to understand the drivers of pest population dynamics and improve pest management practices. Here we analyze spatiotemporal drivers of Colorado potato beetle (CPB, *Leptinotarsa decemlineata*) abundance across Wisconsin potato fields using multi-year scouting data (2014–2024) linked with climate and cropping histories. We develop statistical models that account for spatial and temporal correlations, and find these approaches substantially improve fit and prediction. After accounting for spatiotemporal random effects, we find that three predictors increased CPB abundance: cumulative growing degree days (GDD), recent potato intensity in the surrounding landscape, and winter coldest-day temperature. Cumulative GDD and potato intensity are positively associated with abundance, and warmer winter minima (higher coldest-day temperatures) are likewise associated with higher abundance, consistent with improved overwintering survival. Using the best-performing model, we generate a preliminary, statewide risk surface for Wisconsin in order to support regional decision-making. Our results highlight the value of integrating field-level history with landscape context and explicit spatial structure when forecasting pest pressures in agroecosystems.

## Introduction

Crop pests pose significant challenges, with impacts on regional food security, farm economies, and environmental sustainability (Oerke, 2006; Riegler, 2018). To improve pest management outcomes, there is considerable interest in using ecoinformatic approaches to predict pest population dynamics (Fleischer et al., 1999; Rosenheim et al., 2011; Gerovichev et al., 2021). In particular, long-term large-scale spatiotemporal data sets on pest abundance can be leveraged to refine regional management approaches and improve risk-based decision-making for growers (Lacasella et al., 2017). Spatiotemporal abundance datasets have been shown to effectively predict spatial patterns of pest abundance in unsampled areas (Cohen et al., 2022) and to predict future pest pressure in growing regions (Merrill et al., 2015). Furthermore, the identification of key environmental variables and management practices that influence pest abundance patterns can lead to the refinement of pest management strategies (Ulrichs and Hopper, 2008; Schmidt-Jeffris and Nault, 2018; Machekano et al., 2019).

Pest management of the Colorado potato beetle (CPB, *Leptinotarsa decemlineata*) is an ongoing challenge to sustainable potato production across the United States and globally (Alyokhin et al., 2022). Without effective management, CPB populations can cause severe yield losses in potato crops, and their remarkable ability to evolve resistance to insecticides remains a persistent threat (Alyokhin, 2009; Pélissié et al., 2022). The US Midwestern region is a key agricultural region for potato farming, and understanding the factors influencing CPB population dynamics can inform CPB management practices more broadly. Past research has shown that increasing crop rotational frequency and rotational distance is negatively correlated with CPB abundance, as repeated potato planting at or near a focal field site increases the likelihood of beetle persistence, relative abundance, and insecticide resistance development (Sexson and Wyman, 2005; Huseth et al., 2012; Huseth et al., 2015). Similarly, several key temperature variables, including low winter temperatures and warm summer growing conditions, are known to influence the abundance of CPB (Walgenbach and Wyman, 1984; Ferro et al., 1985; Milner et al., 1992).

Our project addresses two critical gaps in prior research on factors that determine the abundance of CPB. First, many earlier studies lacked spatiotemporal modeling, which is essential because CPB dispersal occurs across heterogeneous landscapes and interannual effects are expected to impact abundance. Second, the relative importance of different environmental and land use predictors have not been assessed in a systematic way. In this study, we assembled an integrated spatiotemporal dataset covering CPB abundance, climate, and cropping patterns across potato fields in Wisconsin between 2014 and 2024. Our goals are to identify predictors of CPB abundance, quantify the effects of historical potato planting intensity at both field and landscape scales, explicitly model spatial and temporal autocorrelation in CPB populations using advanced statistical techniques, and contribute toward long-term efforts to generate predictive risk maps for CPB management.

Here we address these gaps by coupling long-term scouting records with climate and cropping histories and by explicitly modeling spatial and hierarchical structure. We test three predictions: (i) recent landscape-level potato intensity exerts a strong positive effect on CPB abundance; (ii) warmer winter minima increase abundance via improved overwinter survival; and (iii) explicitly modeling spatial correlation materially improves fit and predictive performance relative to non-spatial baselines. We also produce a preliminary, statewide risk map for Wisconsin to illustrate management applications. We evaluated three modeling frameworks of increasing complexity: a linear mixed model (LMM), a generalized additive model (GAM), and a spatial mixed-effects model (spaMM), to assess the contribution of hierarchical structure, nonlinear smoothers, and explicit spatial correlation, respectively. This progression allowed us to identify the model that best captured Colorado potato beetle abundance while comparing the relative contributions of spatial and temporal processes.

## Materials and Methods

### Study Area and Data Collection

Through collaboration with professional pest scouting agencies (Pest Pros Inc., a Division of Allied Cooperative, Adams, WI), we have access to long-term (2008-2024) abundance data on Colorado potato beetle (CPB) from more than 1,800 fields in Wisconsin. Spatial coverage of sampling is centered in central and southern Wisconsin (**Figure 1**). Although scouting data span 2008–2024, we restricted analyses to 2014–2024. Pre-2014 Cropland Data Layer (CDL; USDA NASS 2024) classifications were noticeably noisier in our study area, and our potato-history predictors require a stable five-year look-back window; limiting to 2014 onward provided more reliable crop labels for those histories. The data we use include the number of adult beetles and larvae collected from each field in each year. We averaged these numbers to get one CPB value per field-year, which will be our main response variable. Each field also has its location (latitude and longitude), ownership within a farm operation, and the year that CPB abundance was measured.

**Figure 1.**
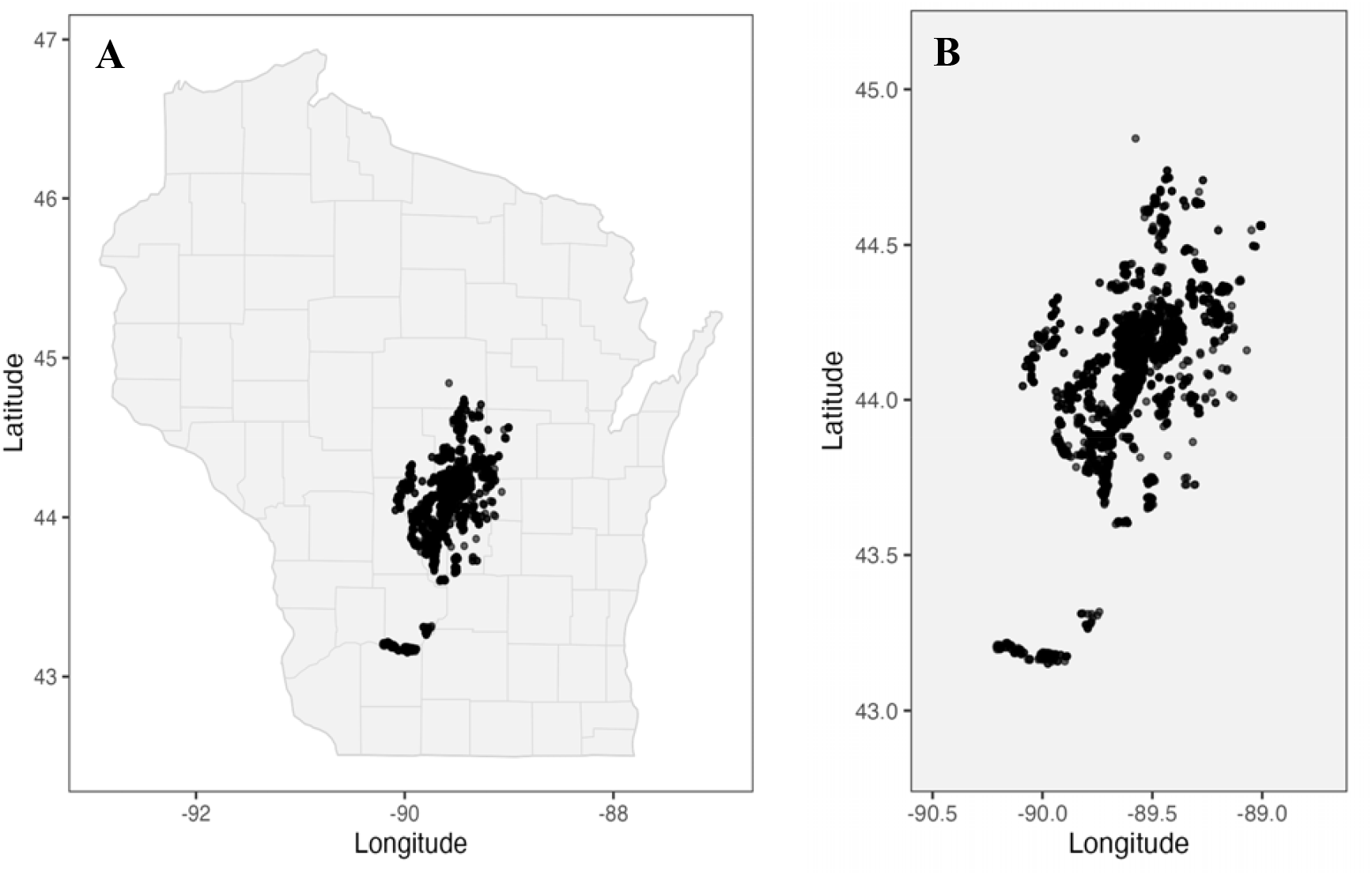
A) Sampling locations across Wisconsin. Each point shows a field included in the scouting dataset (2014–2024), plotted against the state outline. B) Zoomed map of sampling locations, highlighting the core potato production region in central Wisconsin.

Analyses and figures below therefore use 2014–2024 consistently unless otherwise noted. For statewide prediction maps, gridded covariates are fixed to 2021 for comparability across data layers.

### Climate Features

To better understand how weather affects beetle populations, we created several climate variables for each field and year using daily weather data. All of these climate variables were calculated using Python (mainly with the Pandas and NumPy libraries, see associated scripts) based on the daily climate records from GISMETEO (www.gismeteo.ru). These climate data were derived from gridded surfaces with a resolution of 1 km. These were matched based on field location and date. First, we calculated basic values like the average temperature and total rainfall for each year. We also did this for spring (March–May) and summer (June–August), and calculated how much the temperature and rainfall changed throughout the seasons. To capture how long the growing season is, we counted frost-free days (days above 0°C) and noted the date of the last spring frost. If the last frost happens earlier, the beetles might emerge from overwintering diapause earlier and have more time to grow or reproduce. These kinds of changes are especially important because warmer temperatures and fewer frosts may allow beetles to be active earlier and for longer periods. Next, we created variables to represent extreme conditions. We counted how many days in a year were really hot (above 25°C) or really cold (below –25°C), since extreme heat or cold can affect insect survival and development. We also evaluated how often heavy rain occurred during the summer growing season (more than 25 mm in a day), since strong rain can lead to flooding or make the environment colder. This final set of environmental variables provides information on both average conditions and extreme events that can affect beetle populations.

### Crop Features

We compiled crop information using the USDA Cropland Data Layer (CDL; USDA NASS 2024; Li et al., 2024), a 30 m resolution land-cover product. Because field centroids can fall on roads or equipment, we used a circular buffer around each centroid and assigned the modal crop within the buffer to reduce misclassification.

Once annual field crops were assigned, we computed two potato-history metrics. First, a *field-level weighted potato proportion* over a five-year look-back window with integer recency weights 5, 4, 3, 2, 1 for years *t* to *t* − 4:

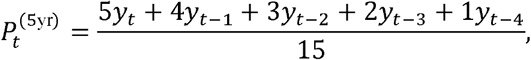

where *y* _*t* −*k*_ ∈ {0,1} indicates potato in year *t* − *k*. This keeps the measure on a 0–1 scale while emphasizing recent plantings.

Second, we quantified *landscape potato intensity* through time (Huseth et al., 2015). For each field-year we drew a 1.5 km buffer (consistent with documented CPB dispersal distances; Follett et al., 1996; Sexson and Wyman, 2005; Boiteau et al., 2008) and computed the proportion of area in potato, again using a five-year, recency-weighted average analogous to 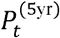. This metric captures nearby host continuity and management intensity through time.

These two features — weighted potato proportion and weighted potato intensity — help us understand whether beetle abundance is dependent on persistence in a field from one year to the next, or dispersal from nearby fields. We calculated these in R using spatial tools that let us measure crop types inside each buffer.

### Assessment of Predictor Variables

We generated ~ 30 climate and crop variables and screened them prior to modeling. We examined pairwise correlations and removed one of any pair with |*r*| > 0.7, favored interpretability, and checked variance inflation (VIF< 5) in pilot linear models. The final set retained 10 predictors spanning seasonal means, extremes, and the two potato-history metrics. All continuous predictors were centered and scaled to zero mean and unit variance prior to fitting; coefficients are therefore on comparable scales, and the same centering/scaling parameters from the training data were used when preparing gridded predictors for statewide prediction.

### Statistical Analysis

We began our analysis by examining the hierarchical structure of the dataset using linear mixed-effects models (LMMs) to test for random effects across farms and years. We first tested whether adding a farm-level, and then a year-level, random intercept improved the model. We then estimated the relative importance of the climate and crop predictors by adding these variables as fixed effects. The LMM can be written as:

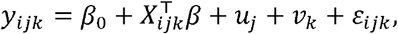

where *y*_*ijk*_ is the CPB abundance for observation *i* in farm *j* and year *k, β*_0_ is the global intercept, *X*_*ijk*_ is the vector of fixed effects (climate and crop predictors), *β* is the vector of associated coefficients, 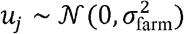 and 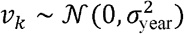 are random effects for *Farm* and *Year*, respectively, and 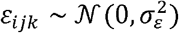 is the residual error.

Next, we used a nonparametric approach to model spatial trends using a generalized additive model (GAM) with smoothing terms over space (*latitude, longitude*) and random smoothers for *Year* and *Farm*. The model includes fixed effects for climate and cropping predictors, and uses thin-plate splines to capture nonlinear spatial and group-level trends. The GAM can be written as:

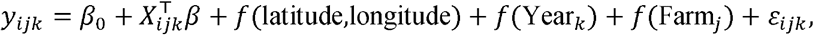

where *f* (·) are thin-plate spline smoothers capturing non-linear spatial or grouping effects.

Third, in order to jointly model spatial correlation and hierarchical structure, we applied a spatial mixed-effects model (spaMM) using the spaMM package (Rousset, 2017). This model includes a Matérn spatial covariance structure over geographic coordinates, as well as random intercepts for *Farm* and *Year*. This approach integrates grouped heterogeneity and continuous spatial dependence into a unified modeling framework. The model can be written as:

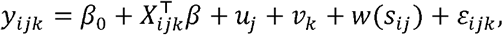

where *w* (*s*_*ij*_) is a spatial random effect governed by a Matérn covariance structure:

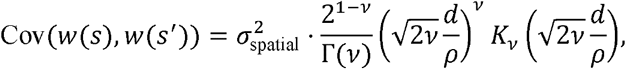

where *d* is the Euclidean distance between coordinates *s* and *s*′, *ρ* is the spatial range, *v* is the smoothness parameter, and *K*_*v*_ is the modified Bessel function of the second kind.

To evaluate the relative performance of the three modeling frameworks, we compared their goodness-of-fit and predictive power using the explained variance (*R*^2^), the mean-squared error (MSE), and Akaike information criterion (AIC).

Unless otherwise noted, we compared models using AIC, marginal and conditional *R*^2^, and MSE computed on the fitted data. Because AIC penalizes complexity and conditional *R*^2^ reflects variance explained including random effects, we rely on the joint pattern (lowest AIC, lowest MSE, highest conditional *R*^2^) to judge overall performance, and we treat the MSE as an in□sample diagnostic rather than a predictive error estimate. All analyses were conducted in R (R Core Team, 2024). LMMs used lme4 (Bates et al., 2015), GAMs used mgcv (Wood, 2017), and spatial mixed-effects models were fit using spaMM (Rousset, 2017).

## Results

### Linear Mixed-effects Model (LMM)

To quantify fixed effects, we fit a full LMM with selected climate and crop predictors. An initial likelihood ratio test comparing a model with a farm-level random intercept to a null model showed strong improvement (ΔAIC = 376, *χ*^2^ = 378.2, *p*< 0.001), indicating substantial variation in CPB abundance across farms (**Table 1**). Extending the model by adding a year-level random intercept further improved fit (ΔAIC = 545, *χ*^2^ = 546.9, *p* < 0.001). These results confirm that both farm- and year-level random effects are important sources of variability— likely reflecting differences in management practices, environmental conditions, and sampling intensity.

**Table 1.** Likelihood ratio tests (LRTs) for random effects in linear mixed-effects.

| Model comparison | df | AIC | logLik | Chisq | p-value |
| --- | --- | --- | --- | --- | --- |
| Null (intercept only) vs. + Farm RE | 1 | 12525 | -6259.4 | 378.18 | $< 2.2 \times 10^{-16}$ |
| Farm RE vs. + Year RE | 1 | 11980 | -5985.9 | 546.92 | $< 2.2 \times 10^{-16}$ |

The full LMM achieved a marginal *R*^2^ = 0.054 (explained variance from fixed effects alone) and a conditional *R*^2^ = 0.444 (total variance explained, including fixed and random effects). The MSE was 2.04, and AIC = 11941.4, providing a baseline for comparison. Two variables were statistically significant and ecologically meaningful (see **Table 2**): cumulative GDD (estimate =0.085, *t* = 2.70) and weighted potato intensity within 1.5 km (estimate = 0.286, *t* = 8.38), indicating higher CPB abundance with greater seasonal heat and in potato-dense landscapes.

**Table 2.** Fixed effect estimates with 95% Wald confidence intervals.

| Term | Estimate | Std. Error | t value | 95% CI Lower | 95% CI Upper |
| --- | --- | --- | --- | --- | --- |
| Intercept | 3.506 | 0.279 | 12.58 | 2.960 | 4.052 |
| GDD | -0.001 | 0.033 | -0.04 | -0.065 | 0.062 |
| Cumulative GDD | 0.085 | 0.031 | 2.70 | 0.023 | 0.147 |
| Weighted Intensity | 0.286 | 0.034 | 8.38 | 0.219 | 0.353 |
| Weighted Proportion | 0.053 | 0.028 | 1.89 | -0.002 | 0.109 |
| Summer Avg. Temp | -0.075 | 0.063 | -1.19 | -0.198 | 0.048 |
| Summer Avg. Precipitation | -0.027 | 0.032 | -0.84 | -0.091 | 0.036 |
| Heavy Rainfall Days | -0.029 | 0.038 | -0.76 | -0.103 | 0.045 |
| Hottest Summer Temp | 0.033 | 0.062 | 0.53 | -0.089 | 0.155 |
| Coldest Winter Temp | 0.082 | 0.067 | 1.22 | -0.049 | 0.213 |
| Winter Extreme Cold Days | 0.026 | 0.057 | 0.45 | -0.086 | 0.138 |
| Winter Warm Day Count | -0.059 | 0.058 | -1.02 | -0.172 | 0.054 |

### Generalized Additive Model (GAM)

We next fit a GAM with the same climate and crop predictors. This model achieved an adjusted marginal *R*^2^ =0.344 and explained 36.7% of the deviance, indicating stronger fixed-effect fit than the LMM. Both the MSE = 1.94 and the AIC = 11604.5 improved. Four predictors were significant (see **Table 3**): cumulative GDD (estimate = 0.094, *p* = 0.0025), weighted intensity (estimate = 0.225, *p*< 0.001), heavy summer rainfall days (estimate = −0.086, *p* = 0.024), and winter coldest-day temperature (estimate = 0.182, *p* = 0.008). Weighted proportion was not significant after accounting for these effects. For the smoothing terms, the effective degrees of freedom were highly significant for space, year, and farm (**Table 4**), indicating nonlinear spatial structure and grouping heterogeneity.

**Table 3.**
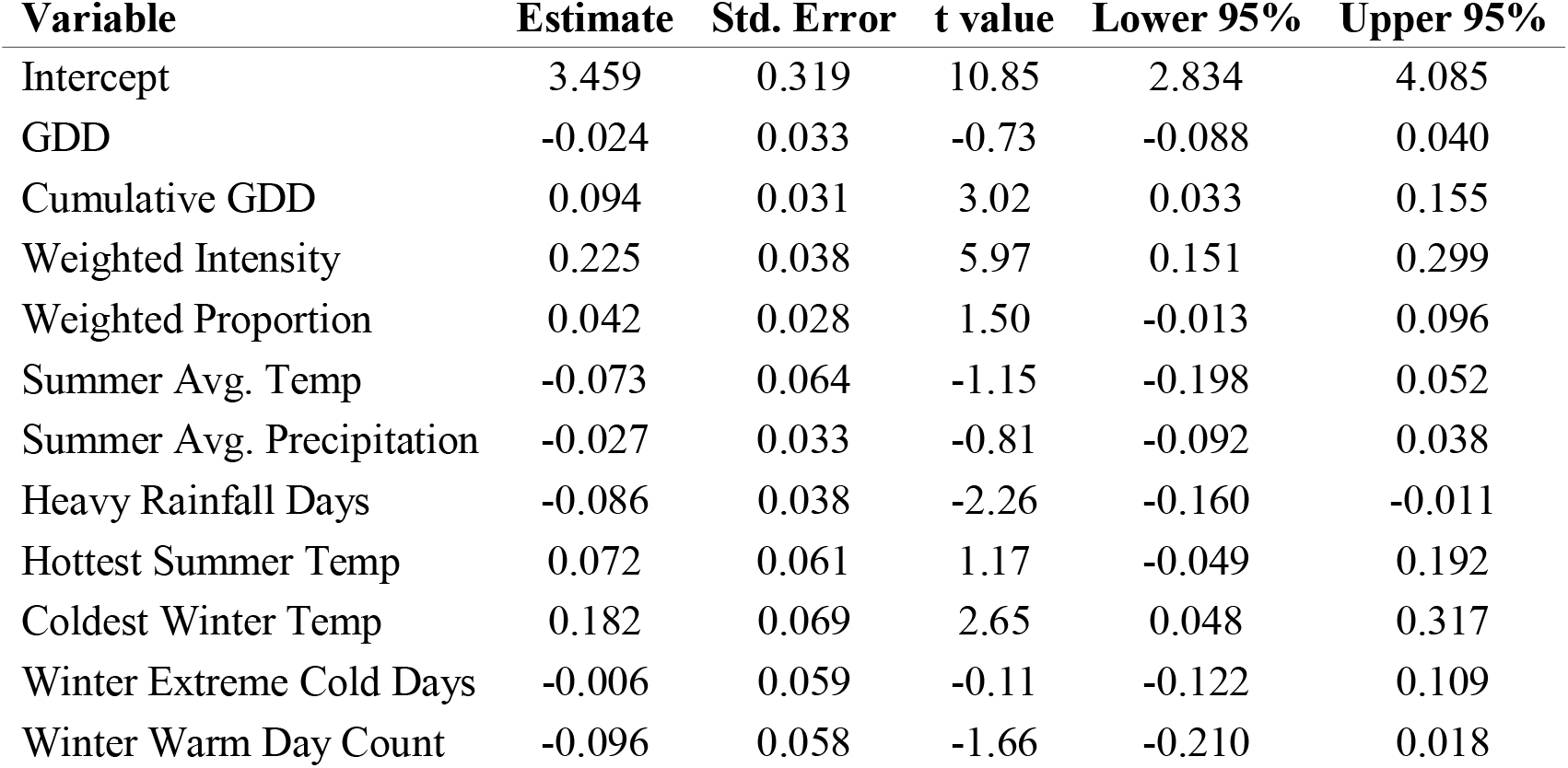
Fixed effects (with 95% CIs) from the generalized additive model.

**Table 4.** Significance of smooth terms in the generalized additive model.

| Smooth Term | EDF | F-statistic | p-value |
| --- | --- | --- | --- |
| s(lat, lng) | 11.12 | 11.55 | < 0.001 *** |
| s(year) | 9.78 | 54.22 | < 0.001 *** |
| s(farm) | 48.93 | 5.97 | < 0.001 *** |

Additional plots of spatial and random smoothers estimated under the GAM model (**Supporting Information Figure S1**) show the spatial smooth *s* (*lat, lng*), as well as the random effects of year and farm. We emphasize that these diagnostics come from the *GAM*, whereas our main spatial inference in the text relies on the *spaMM* model that explicitly incorporates Matérn spatial correlation.

### Spatial Mixed-effects Model (spaMM)

We next fit a spaMM with the same climate and crop predictors. This model achieved the highest total explanatory power 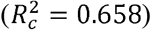 among all models tested, with marginal *R*^2^ = 0.188 Both the MSE = 1.795 and the AIC = 11548.3 improved further. The same four predictors highlighted by the GAM were significant under spaMM (see **Table 5**): cumulative GDD (estimate = 0.101, *t* = 3.28), weighted intensity (0.183, *t* =.3.92), heavy rainfall days (−0.081, *t* = −2.13), and winter coldest-day temperature (0.149, *t* = 2.13). The estimated Matérn parameters (*v* = 0.394, *ρ* = 0.809; **Table 6**) indicate a relatively rough spatial surface (smaller *v* implies less smoothness) and a practical correlation range on the order of tens of kilometers. Because the model uses (lat,lng) in degrees, *ρ* = 0.809 corresponds to roughly 80–90 km at 44^0^N (1^0^ latitude ≈ 111 km; 1^0^ longitude ≈ 80 km). Thus, fields within ~ 80–90 km exhibit appreciably correlated deviations after accounting for covariates, with correlation decaying beyond this scale. The random-effect variances further show a substantial spatial component relative to year and farm, reinforcing the importance of explicit spatial covariance in this system.

**Table 5.**
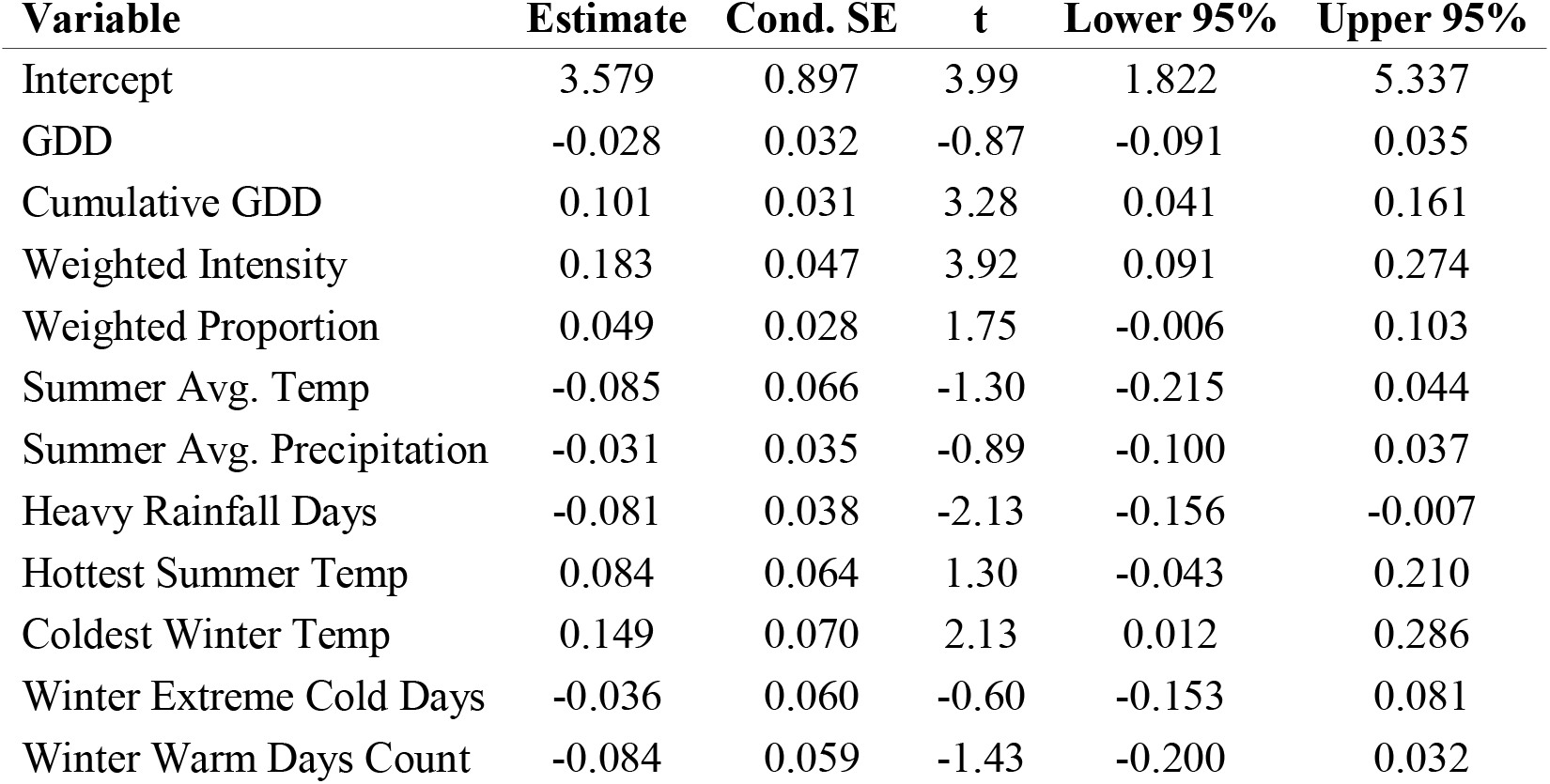
Fixed effects (with 95% CIs) from the spatial mixed-effects model (spaMM); Matérn + Year + Farm.

**Table 6.** Spatial and random-effect parameters from the spaMM model, showing spaMM with lowest AIC/MSE and highest conditional *R*^2^.

| Parameter | Estimate |
| --- | --- |
| Matérn smoothness ($\nu$) | 0.394 |
| Matérn range ($\rho$) | 0.809 |
| Variance (spatial: lat+lng) | 1.503 |
| Variance (year) | 0.625 |
| Variance (farm) | 0.477 |
| Residual variance ($\phi$) | 1.894 |

The residual spatial correlation surface (Matérn effect; **Figure 2**) visualizes residual spatial structure after controlling for climate, cropping history, and year/farm effects. Warmer colors indicate areas where observed CPB abundance is higher than expected from covariates (positive deviations), while cooler colors indicate lower-than-expected abundance (negative deviations). This map confirms local spatial dependence—providing visual justification for the Matérn term—and reveals a broad southwest–northeast gradient consistent with the estimated covariate effects and variance components.

**Figure 2.**
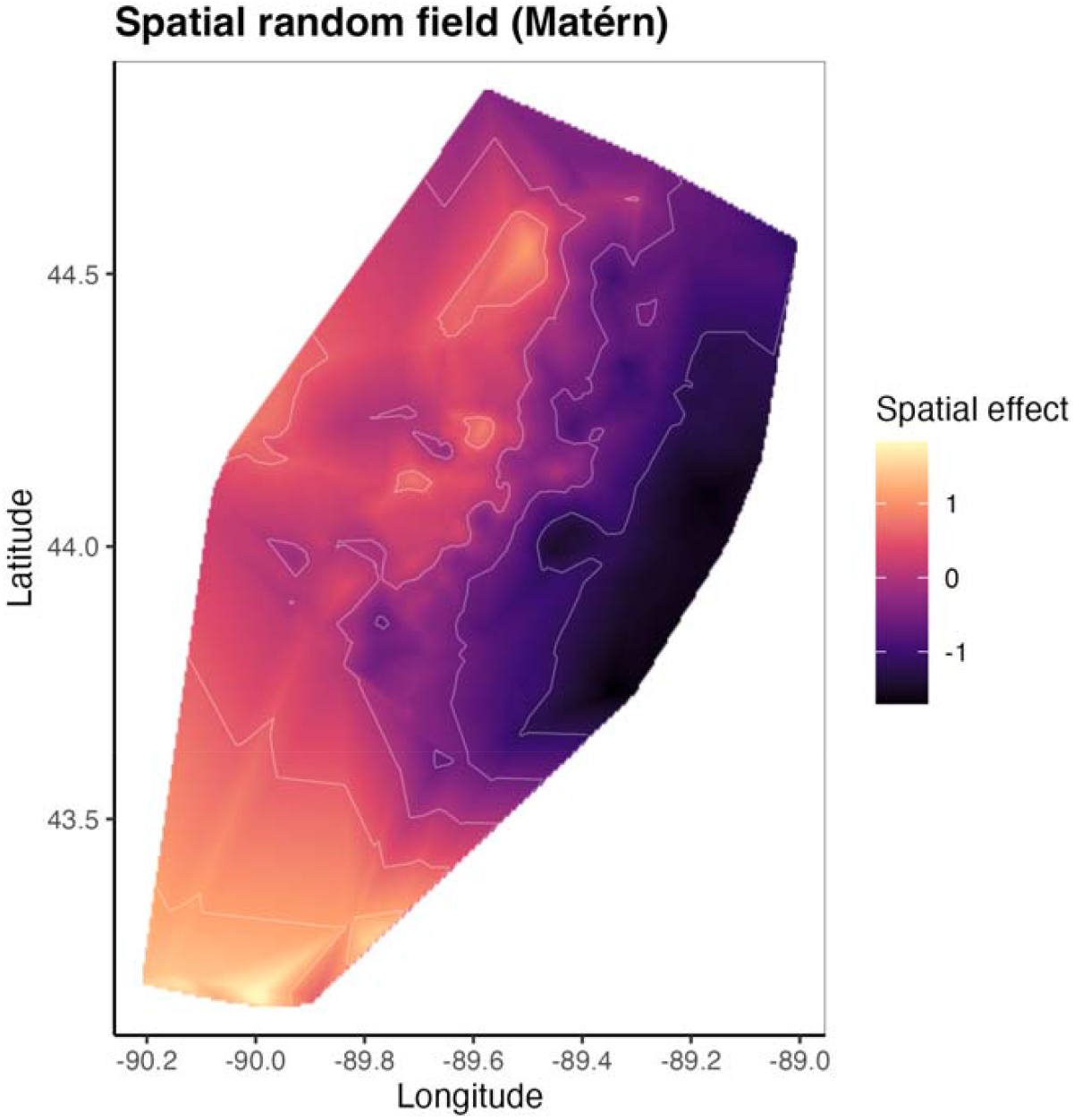
Residual spatial correlation surface (Matérn effect), shown for the core potato production region in central Wisconsin. Warmer colors indicate positive residual deviations (higher-than-expected abundance given covariates); cooler colors indicate negative residual deviations.

### Model Comparison and Interpretation

All three modeling approaches underscore the importance of incorporating both spatial and temporal structure in understanding CPB abundance (see **Table 7**). The spaMM model achieved the lowest AIC and MSE and the highest total explained variation (conditional *R*^2^ = 0.344), demonstrating the value of explicit spatial covariance. The GAM provided the best marginal explanatory power (adjusted *R*^2^ = 0.344), suggesting that flexible smoothers capture nonlinear effects—particularly for space and time. Partial-effect curves from the spaMM fit illustrate key relationships: abundance increases with cumulative GDD and with recent landscape potato intensity, decreases with more heavy summer rainfall days, and increases with warmer winter minima (**Figure 3**). Residual diagnostics indicate approximate homoscedasticity with mild right□tail deviations from normality, consistent with a few high□abundance fields (**Supporting Information Figure S2**).

**Table 7.**
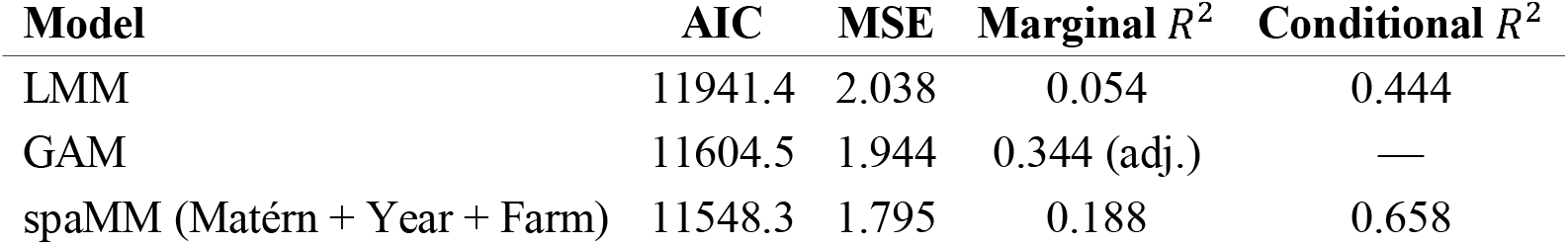
Comparison of spatiotemporal statistical models of Colorado potato beetle abundance in Wisconsin.

| Model | AIC | MSE | Marginal $R^2$ | Conditional $R^2$ |
| --- | --- | --- | --- | --- |
| LMM | 11941.4 | 2.038 | 0.054 | 0.444 |
| GAM | 11604.5 | 1.944 | 0.344 (adj.) | — |
| spaMM (Matérn + Year + Farm) | 11548.3 | 1.795 | 0.188 | 0.658 |

**Figure 3.**
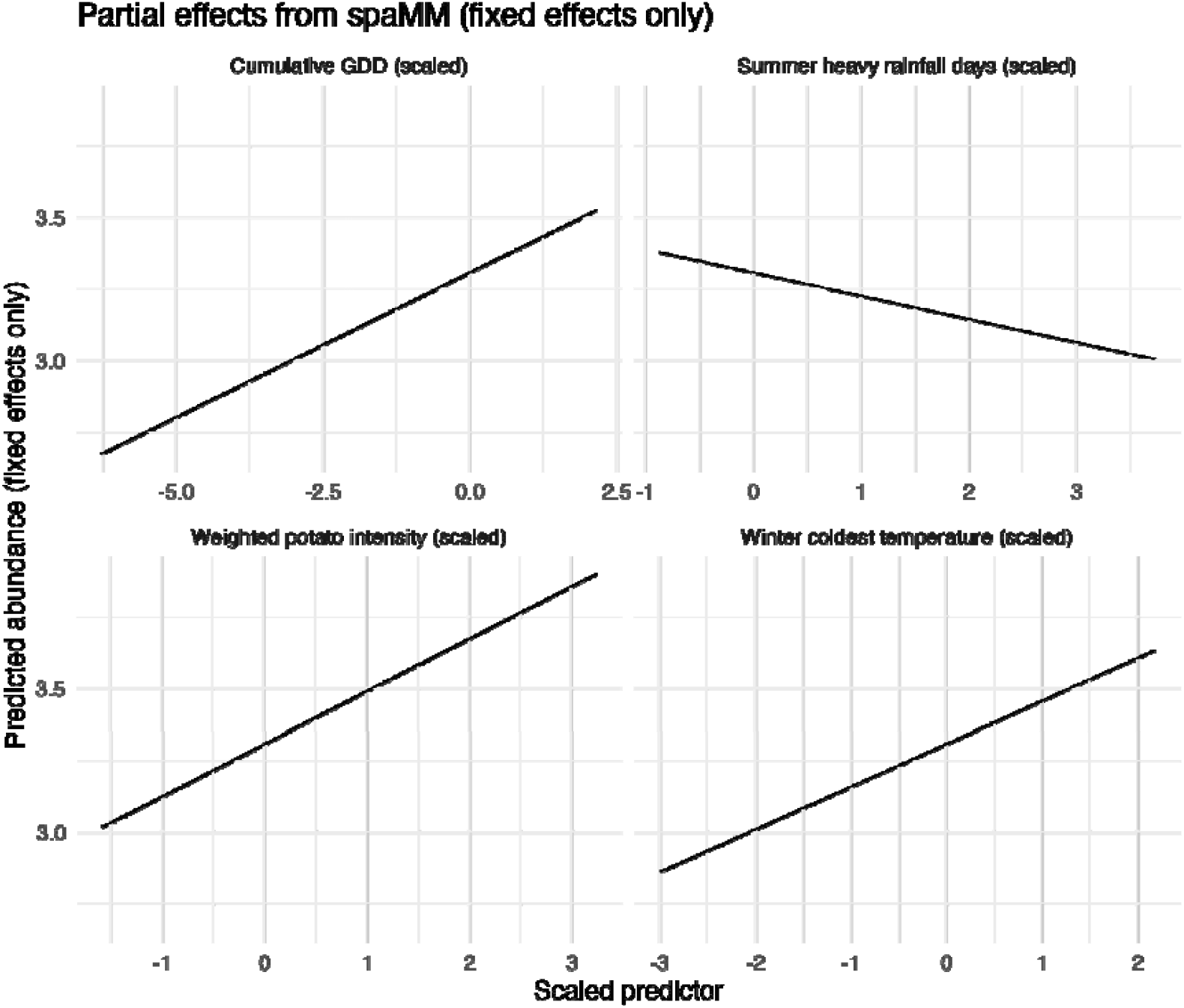
Partial effects of significant predictors from the spatial mixed-effects model (spaMM), showing marginal relationships between predictors and Colorado potato beetle abundance.

### Predicted Risk Surface

We generated statewide predictions of CPB abundance on a 10 km grid using the final spatial mixed model (spaMM; Matérn + Year), setting *farm* random effects to their population mean (i.e., omitted for out-of-sample prediction). Predictor variables were standardized using the means and standard deviations from the modeling dataset to ensure consistency between fitting and prediction. The prediction year was fixed to 2021 as a representative year. Predicted abundances were linearly scaled to a 0–1 index for interpretability, where darker colors indicate lower relative risk (near 0) and lighter/yellow colors indicate higher relative risk (near 1). The resulting statewide pattern (**Figure 4**) highlights spatial heterogeneity, with elevated risk concentrated in regions of greater recent potato intensity and milder winter minima. Scenario maps (potato intensity × 0.7 and × 1.5) are provided in the (**Supporting Information Figure S2**) to illustrate sensitivity to this key driver. Uncertainty in predictions is greater near the range margins and in regions with limited training data, so the map should be viewed as a relative risk index rather than an absolute forecast. Because farm-level random effects were set to their population mean, the surface reflects landscape and climate structure rather than farm-specific baselines. These choices ensure the map is most useful for regional planning and scouting strategies, rather than precise field-level forecasts.

**Figure 4.**
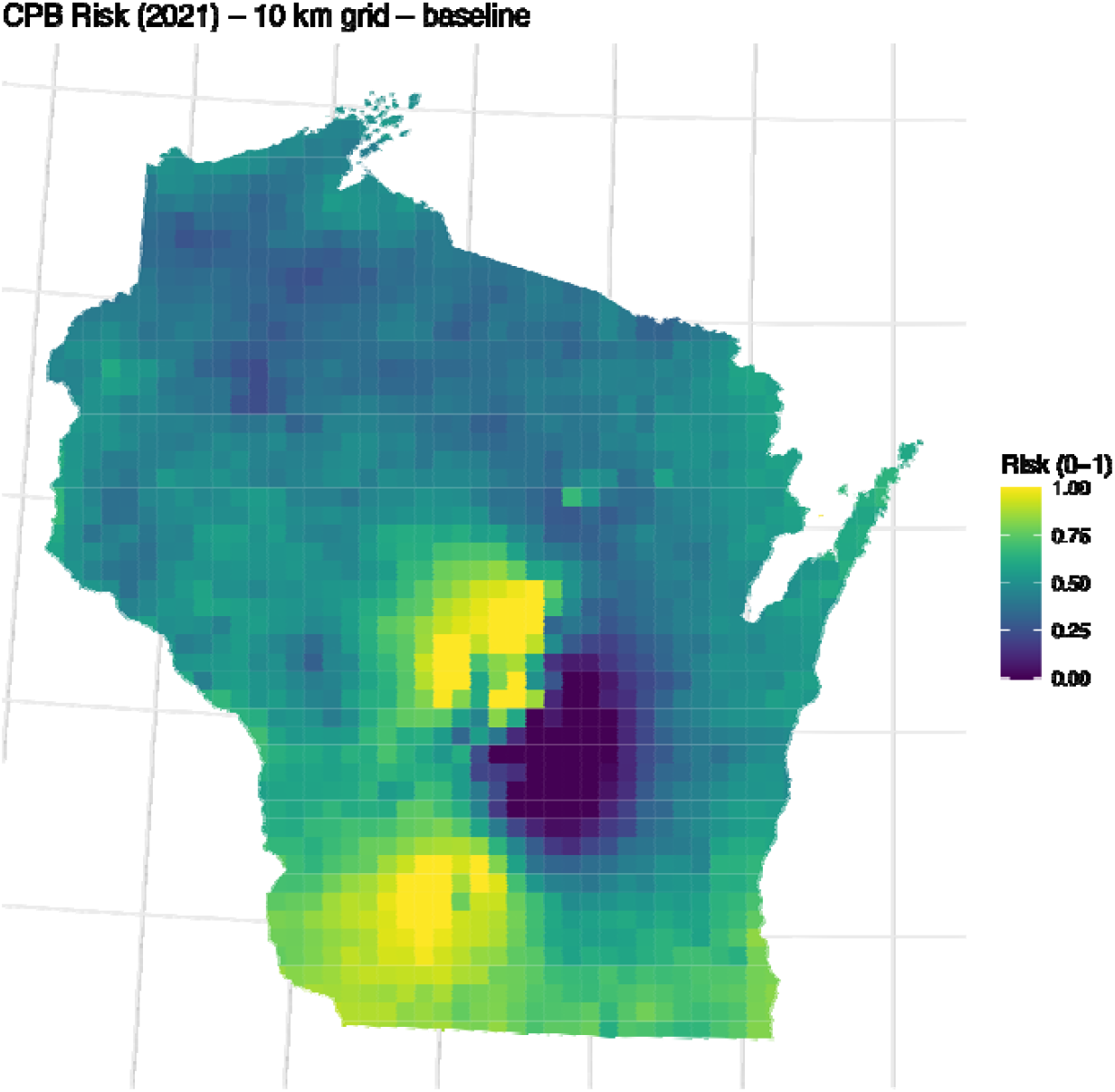
Statewide CPB risk surface for Wisconsin predicted on a 10 km grid using the spaMM model (Matérn spatial field + year random intercept). Predictions omit farm-level random effect, fix the prediction year to 2021, and are rescaled to a 0–1 risk index. Darker colors indicate lower relative risk (near 0), and lighter/yellow colors indicate higher relative risk (near 1).

## Discussion

Large-scale datasets of pest abundance are emerging and are increasingly used to support precision agriculture and decision-making (Abd El-Ghany et al., 2020). However, robust inference and forecasting require models that account for spatial and temporal dependence in pest populations (e.g., Schmidt-Jeffris and Nault, 2018). By applying three different statistical frameworks—linear mixed models (LMM), generalized additive models (GAM), and a spatial mixed-effects model (spaMM)—we were able to evaluate not only the role of fixed effects but also the contribution of spatial and temporal structure to overall variation in beetle populations. Our analysis shows that both spatial and temporal correlations exert measurable influences on Colorado potato beetle (CPB) abundance across Wisconsin. Colorado potato beetle abundance across Wisconsin is shaped by both environmental forcing and landscape history, cumulative growing degree days, recent potato intensity within 1.5 km, and warmer winter minima were positively associated with abundance, while heavy summer rainfall days showed a negative association. Explicitly modeling spatial covariance (spaMM with a Matérn field) yielded the best fit and lowest error, indicating that neighboring fields share residual signal beyond measured covariates. This improves our ability to forecast risk of CPB pest pressure in potato agroecosystems.

### Key Predictors of CPB Abundance

Across models, several predictors consistently emerged as influential. Cumulative growing degree days (cumGDD) were positively associated with beetle abundance, reflecting the importance of accumulated seasonal warmth for insect development and activity (Ferro et al, 1985). Potato cropping intensity in the surrounding landscape also showed a strong positive effect, confirming that local landscape history shapes beetle pressure through overwintering and short-distance dispersal (Boiteau et al., 2008; Huseth et al., 2015). Warmer winter minima (higher coldest winter temperatures) were similarly linked to greater abundance, consistent with the idea that milder winters allow more beetles to survive diapause (Milner et al., 1992; Hiiesaar et al., 2006).

At the same time, certain weather extremes appeared to reduce beetle numbers. In particular, an increase in heavy summer rainfall days was negatively associated with abundance. This is ecologically plausible, as intense rainfall events can increase larval mortality or disrupt beetle development (Chen et al., 2019). Other variables, such as short-term GDD, average seasonal temperature, and extreme cold days, were not as consistently significant once the core predictors were accounted for. These patterns suggest that CPB populations are most strongly shaped by longer-term climatic windows and predictable access to potato hostplants, rather than short-term fluctuations alone. While many of these factors have previously been linked to CPB abundance, the influence of spatial and temporal effects on the relative importance of different factors was not known.

### Importance of Spatial and Temporal Structure

Longterm data on insect abundance have been increasingly important in dissecting the factors that impact variability and have consequences for agricultural management (Paredes et al., 2022). In our study, model comparisons highlighted the value of explicitly accounting for spatial and temporal structure (**Table 1** and **Table 7**). The spaMM model, which included both Matérn spatial covariance and random intercepts for year and farm, achieved the best overall performance (lowest AIC and MSE, highest conditional *R*^2^). This finding underscores the importance of spatial dependence: fields closer together in space experienced more similar levels of CPB population pressure than would be expected by chance, even after controlling for climate and crop predictors. This spatial signal likely reflects unmeasured drivers such as landscape management practices, soil conditions, or local dispersal pathways. Thus, spatial proximity may serve as a proxy for aggregate effects of local environmental and management variables not captured in the model.

The GAM also provided useful insights, especially in capturing nonlinear trends over space and time. Its higher marginal *R*^2^ suggests that smoothers can explain fixed-effect variance efficiently, though without the same overall explanatory power as the spaMM. The LMM, by contrast, was more limited, explaining only a small portion of the fixed-effect variation but confirming the importance of farm- and year-level heterogeneity.

### Ecological and Management Implications

These results have several implications for pest management. First, they reinforce the role of crop history in structuring and maintaining elevated CPB populations: fields with frequent potato planting, or those embedded in potato-intensive landscapes, face consistently higher beetle pressure, maintaining elevated pest pressure over time (Sexson ad Wyman, 2005). This can also influence insecticide resistance management (Huseth et al., 2015; Crossley et al., 2022), as strong sustained selection on the same population drives resistance evolution. This emphasizes the need for crop rotation strategies, both spatial and temporal, that reduce local host continuity and CPB populations. This aligns with core principles of pest management that emphasize spatial isolation and disruption of host continuity (Schmidt-Jeffris and Nault, 2018, but also see Rosenheim et al., 2022). Second, climatic effects—particularly milder winters and cumulative heat—are likely to exacerbate beetle problems under ongoing climate change (Hiiesaar et al., 2013). Regional warming trends may enable larger overwintering populations and earlier, and perhaps more protracted, seasonal emergence or multiple generations, amplifying pest risks (Walgenback and Wyman, 1984). They will also increase pest pressure in more northern growing regions (Jönsson et al., 2013). Third, the negative impact of extreme weather events may provide some natural suppression, such as cold winters without snow (Milner e al., 1992) or heavy summer rainfall (Chen et al., 2019), though this is unlikely to offset the long-term increases in abundance driven by warming.

From a modeling perspective, the superiority of spaMM demonstrates the necessity of incorporating spatial correlation into regional pest forecasting. Failing to account for spatial dependence risks overestimating predictor effects and underestimating uncertainty. Our findings also suggest that GAMs remain useful exploratory tools for identifying nonlinearities and smooth spatiotemporal patterns, while LMMs provide a simpler baseline for implementing models with hierarchical structure.

### Toward Predictive Risk Mapping

Importantly, this work contributes to the development of statewide risk maps for CPB in Wisconsin. By integrating field-level cropping history, landscape context, and climate predictors into a spatially explicit model, we can generate predictive surfaces of beetle abundance that highlight high-risk regions. The resulting statewide risk surface (**Figure 4** and **Figure S3**) shows elevated risk concentrated in the potato production core, with moderate values extending into surrounding areas where potato is not grown (urban or forested land). Very low values occur in wetter southern and eastern regions, consistent with the negative association between heavy rainfall and beetle abundance. These heterogeneous but interpretable patterns highlight management□relevant hotspots aligned with potato intensity and winter conditions.

Risk-mapping tools can support proactive decision-making by growers and pest management agencies, guiding scouting efforts, rotational planning, and resistance management strategies. Importantly, our results show that risk is not evenly distributed but is concentrated where potato planting is frequent and winters are warmer, a pattern that is becoming, or has already become, more pronounced under recent climate trends. If climatic warming trends continue, and potato production is increasingly concentrated in compact growing regions, CPB abundance will become a more significant problem. These insights can inform both short-term management and long-term strategies for reducing pest economic impacts.

Overall, our study demonstrates the value of combining ecoinformatics with advanced spatiotemporal models to better understand pest population dynamics (Rosenheim et al., 2011). The integration of landscape history, climate extremes, and spatial correlation provides a more complete picture of CPB abundance, and highlights pathways for both improved forecasting and more sustainable management of this persistent agricultural pest.

## Acknowledgments

The authors wish to thank Pest Pros Inc. and the cooperating membership of the Wisconsin Potato and Vegetable Growers Association for access to crop data.

## Data Availability Statement

All data and scripts are available on GitHub at https://github.com/drnursultan/Spatiotemporal_CPB_Modeling.

