## Supporting Information for "Regional analysis of spatiotemporal trends in Colorado potato beetle abundance"

| 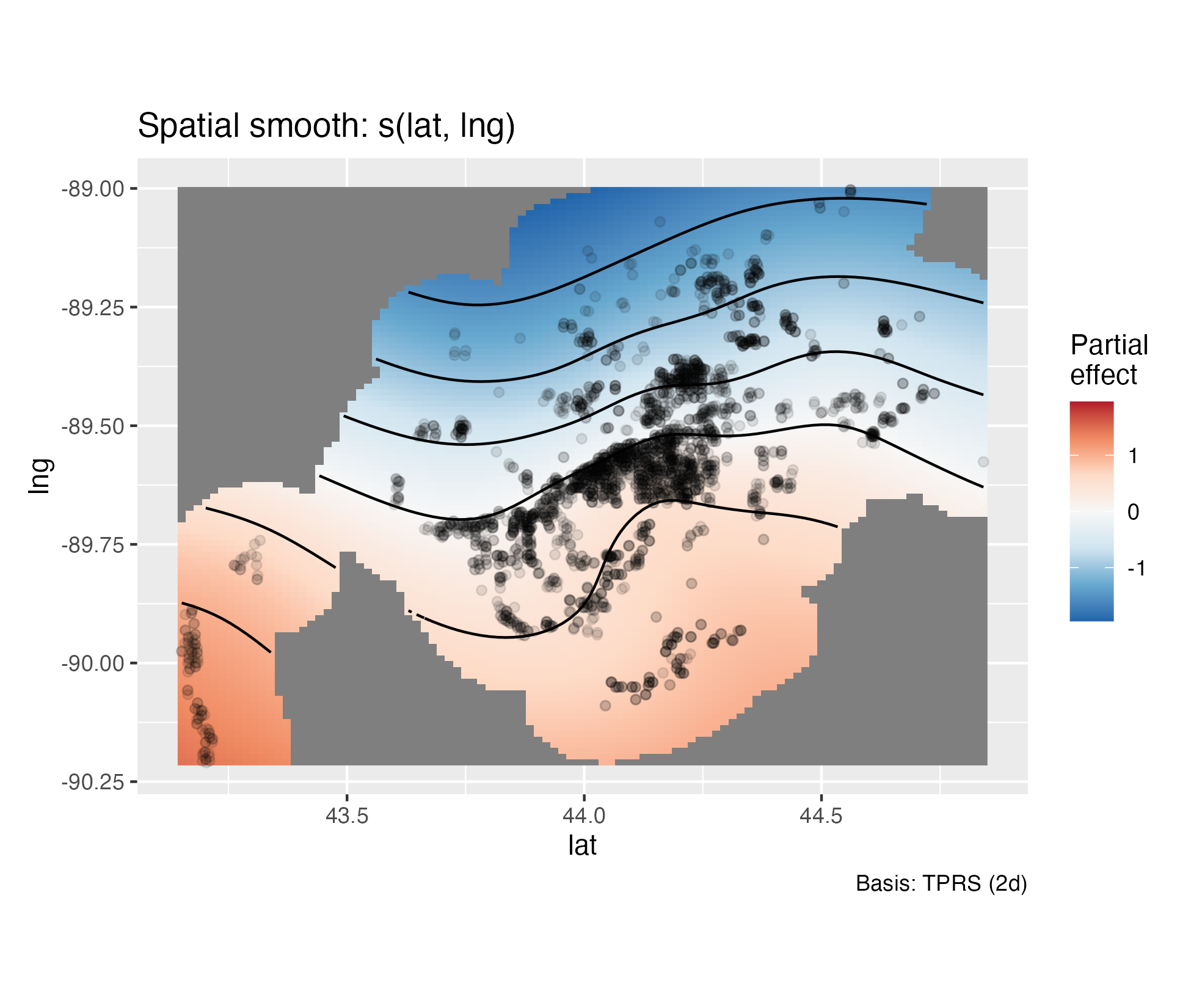 | 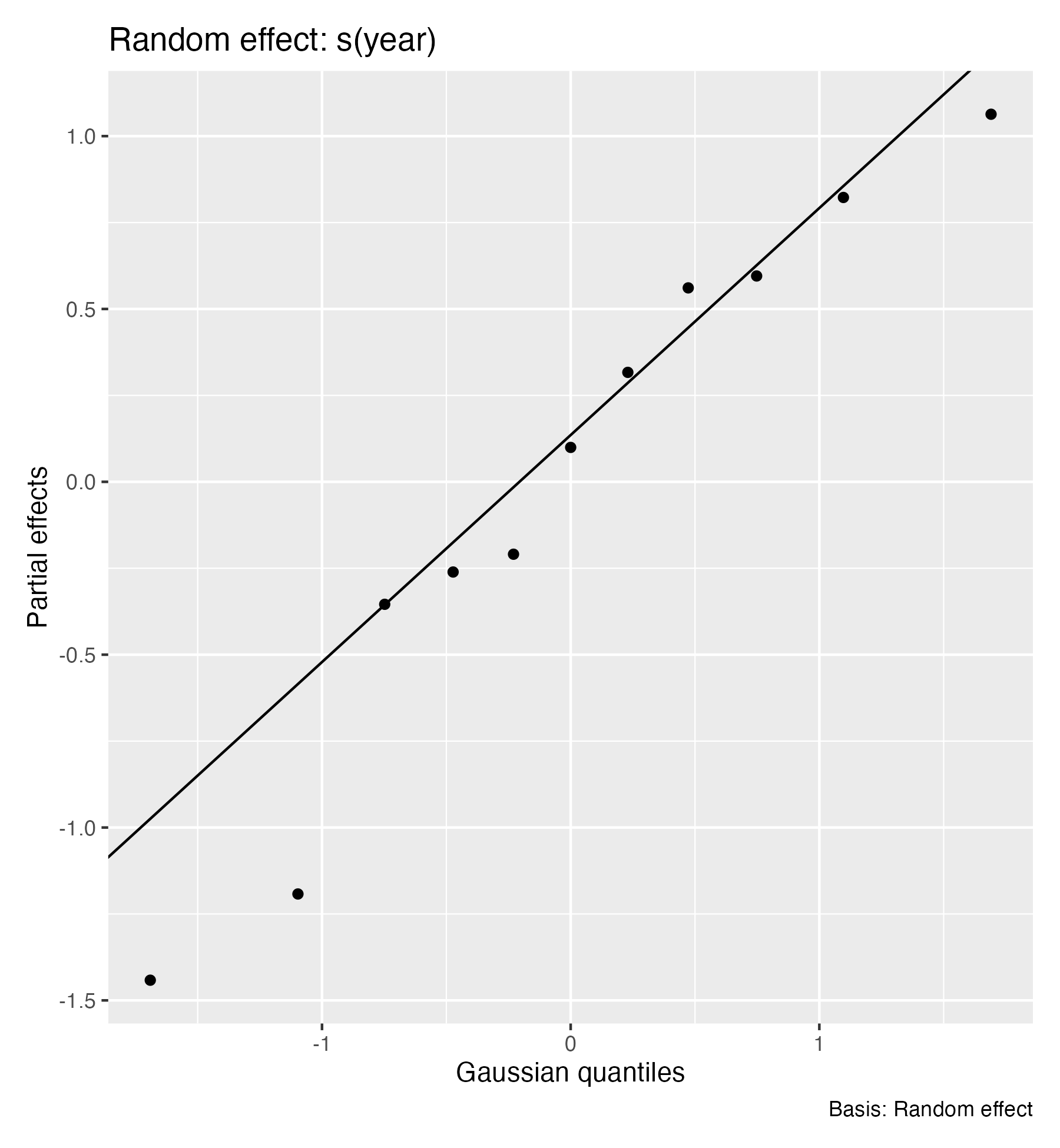 | 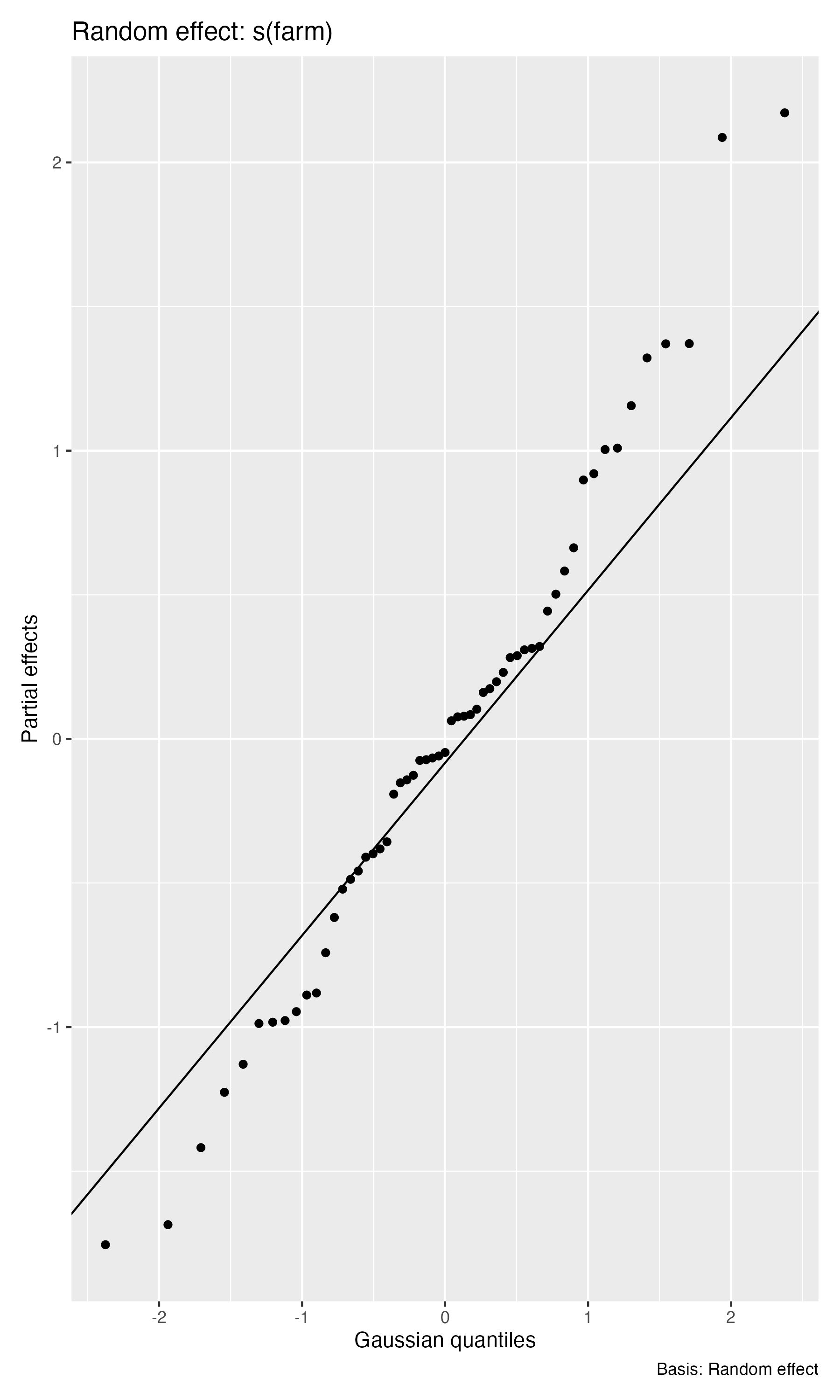 |
| --- | --- | --- |

**Figure S1**. Generalized additive model (GAM) diagnostics. Partial effects showing the spatial smoother and the random-effect smoothers for year and farm.

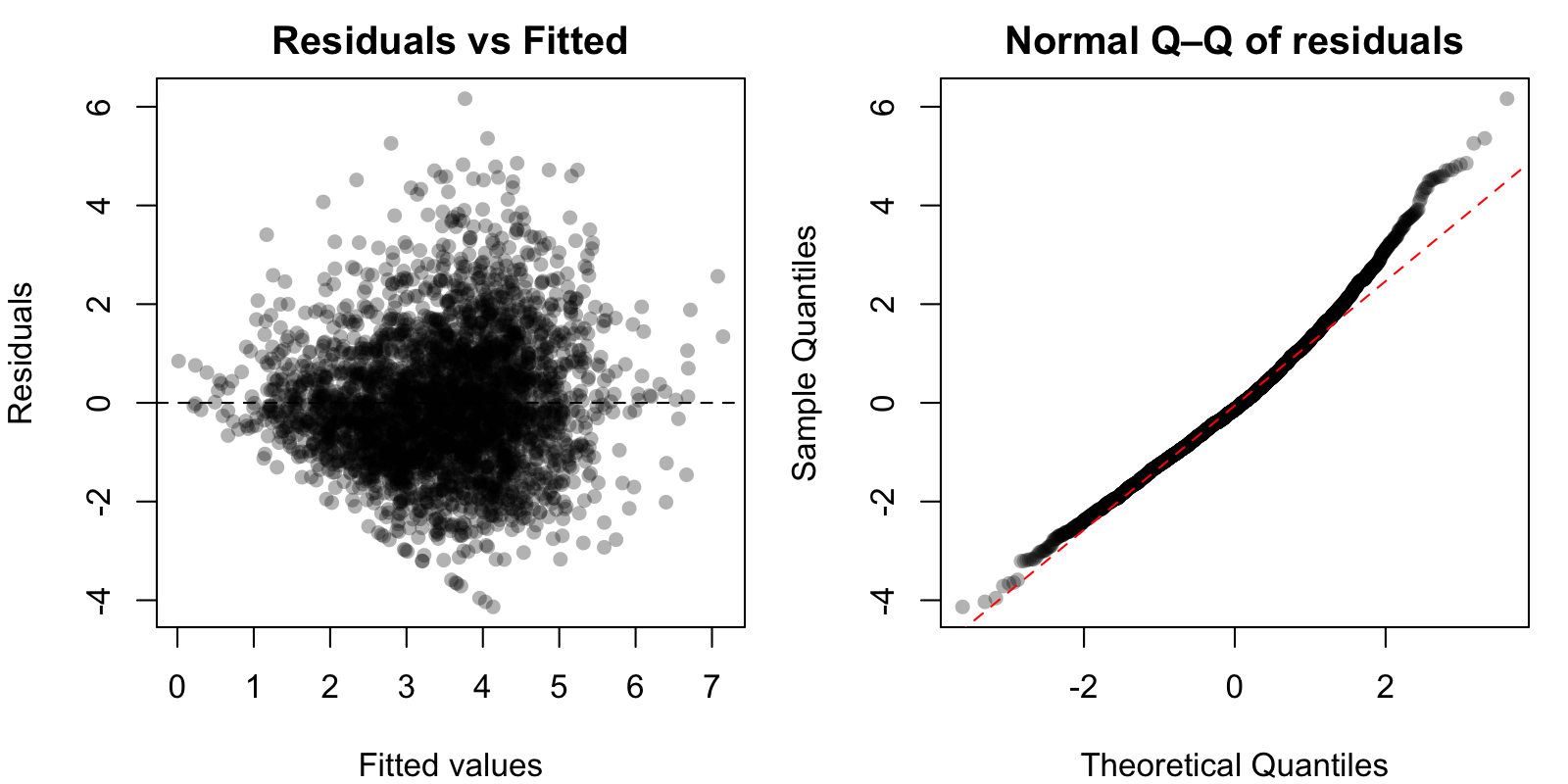

**Figure S2**. Residual diagnostics from the spaMM model, including quantile–quantile plots, residual vs. fitted scatterplots, and residuals over space.

| 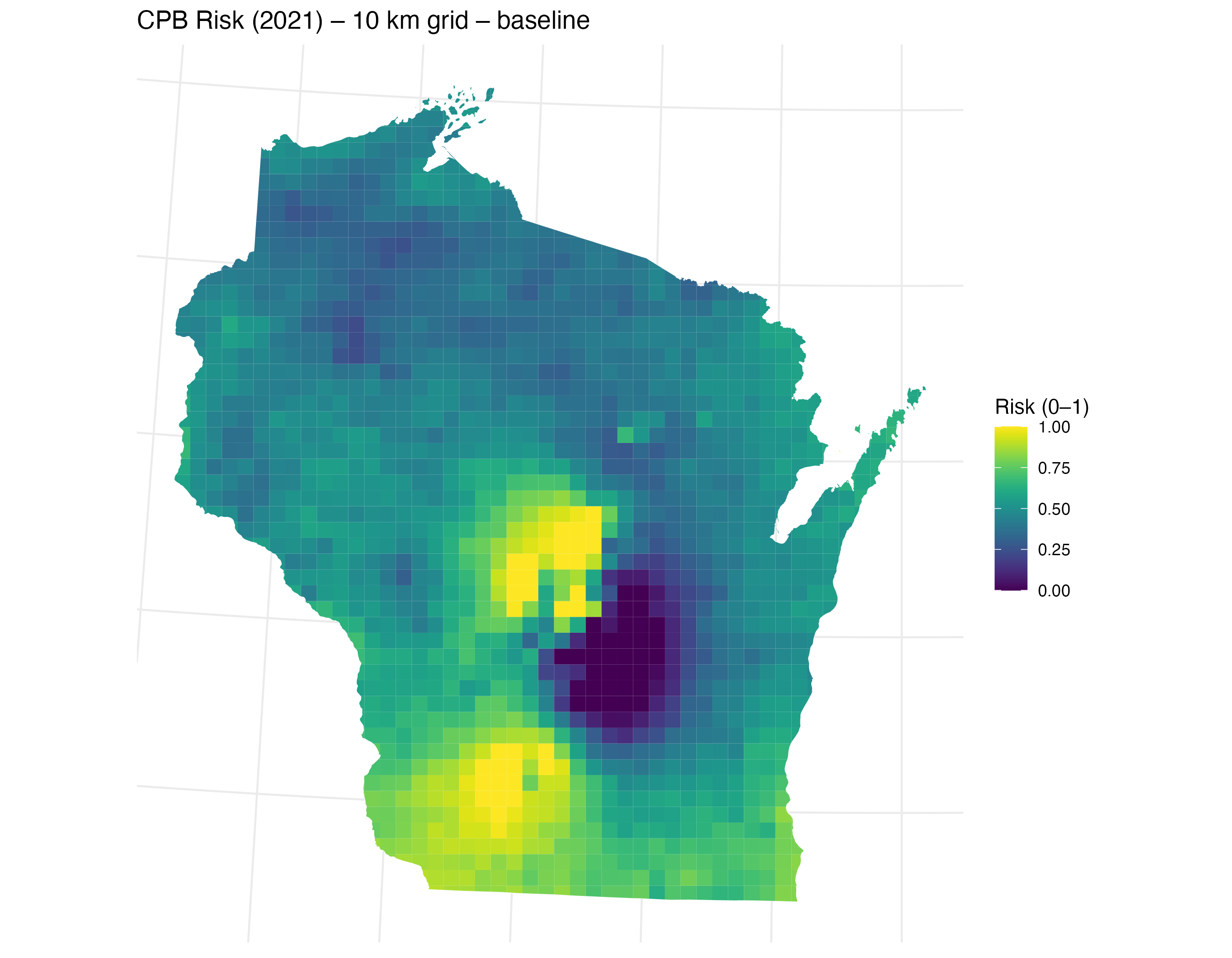 | 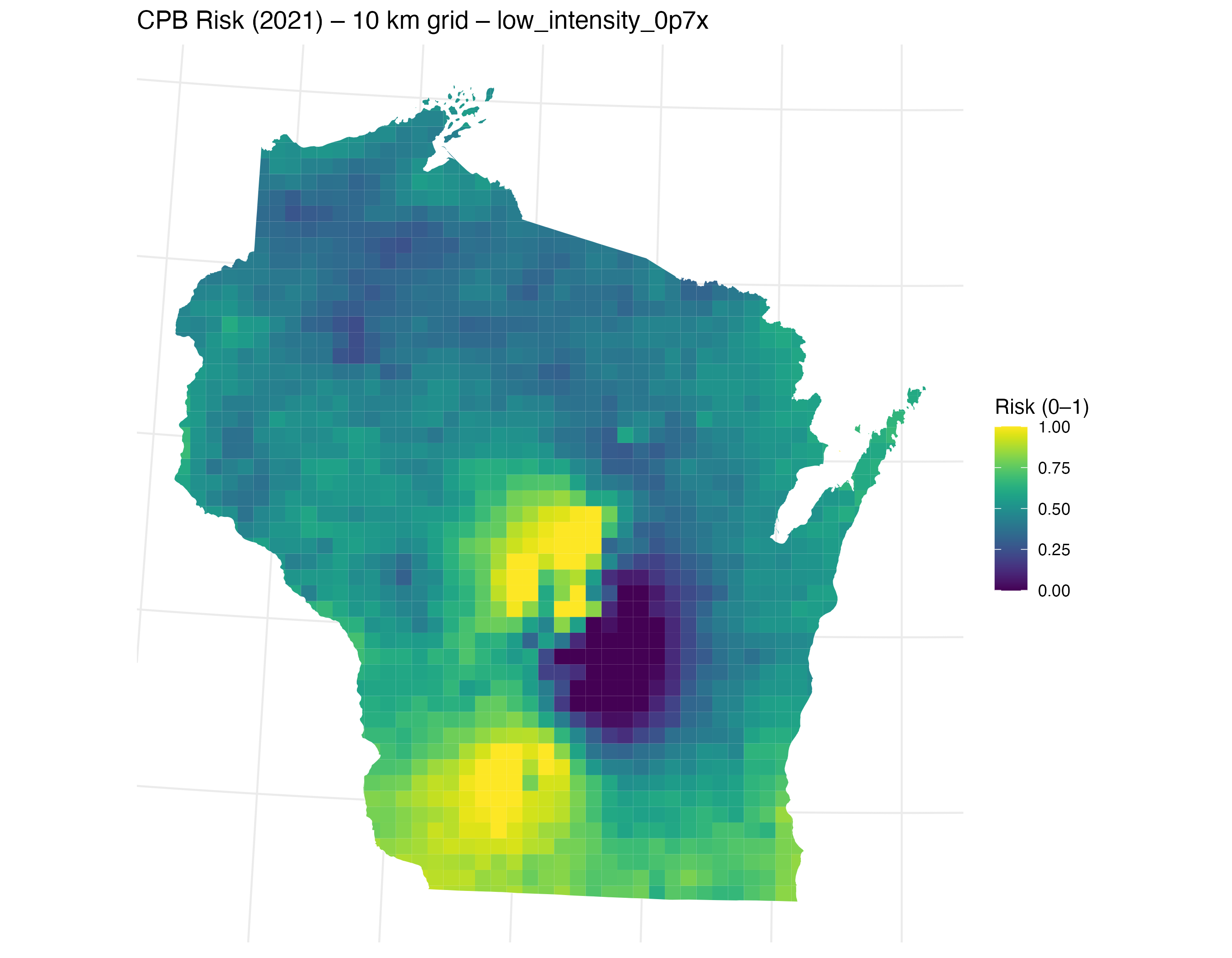 | 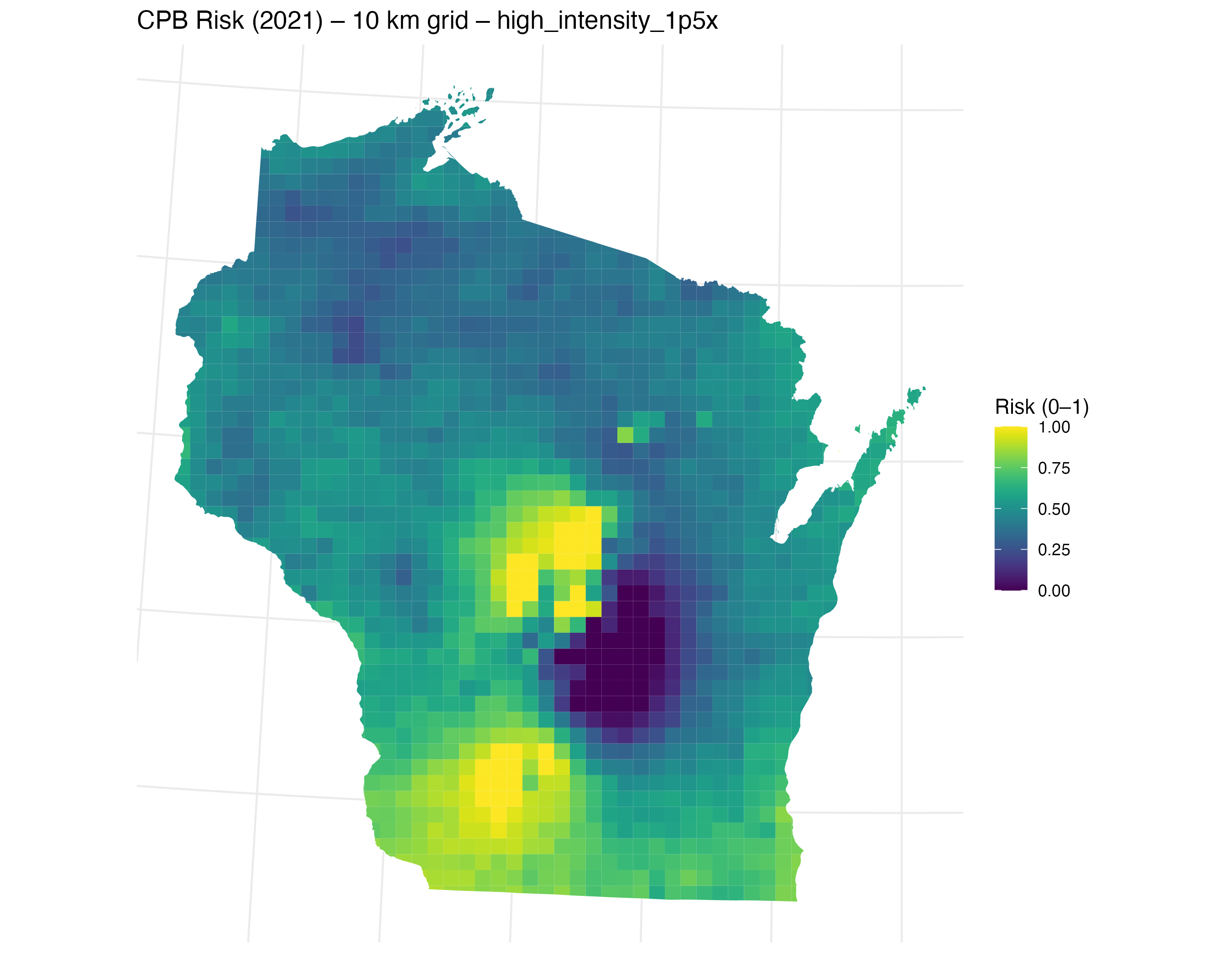 |
| --- | --- | --- |

**Figure S3**. Scenario maps from the spaMM model showing the statewide CPB risk surface when the landscape potato intensity within 1.5 km is held at (a) baseline, (b) reduced by 30%, and (c) increased by 50%. Other predictors fixed as in the baseline map; year fixed to 2021; farm random effects omitted for out-of-sample generalization. Risk rescaled to a 0–1 index.
